# Colorimetric Detector for Hydrogen Sulfide Detection in Industrial Environments Based on Silver Nanoparticles Synthesized in Streptomyces Bacteria

**DOI:** 10.1101/2023.01.25.525489

**Authors:** Nika Souri, Maedeh Abbaszadeh, Sadegh Ghorbanzadeh

## Abstract

Hydrogen sulfide is a colorless, foul-odor pollutant with corrosive properties. This gas is one of the important pollutants released from food factories, fertilizer manufacturing factories, sewage treatment plants, and well drilling. Considering the dangers of being exposed to hydrogen sulfide gas, this article provides a cost-effective, simple, life-saving efficient solution. In this study, Streptomyces bacterium was used for biological production of silver nanoparticles, due to its resistance to the antimicrobial properties of silver. The synthesis and the size range of nanoparticles were confirmed using validation methods. For preparing biocomposite that entraps biological agents, microencapsulation of bacteria was adopted using gellan gum and alginate. In order to validate the final product, the viscosity and porosity of the biocomposite were determined. These tests aimed to investigate the mechanical properties affecting the behaviour and the response time of the final detector. In the end, the detector was exposed to hydrogen sulfide gas, and the color changes of the detector were recorded by time. According to the results, the control sample remained unchanged in all concentrations. However, the color of the test sample, changed faster with the increase in the input gas concentration. Moreover, no color change was observed in concentrations less than 20 mg/l. Therefore, the designed biodetector can be used for qualitative and semi-quantitative measurement of Hydrogen sulfide, in concentrations higher than 20 mg/l.

## I. Introduction

Hydrogen sulfide (H_2_S) is a natural compound existing in many physical and biological systems. This compound is a very toxic, colorless, flammable gas, with a rotten egg-like smell in high concentrations, and heavier than air, dangerous for workers, equipment, and the environment. Anyone who is exposed to H_2_S gas in their workplace are responsible to protect themselves and others from the deadly effects of this gas. Hydrogen sulfide concentration of 100ppm can cause lung blockage and increase heart disease risks. A concentration of 300-500 ppm causes lung edema, and a concentration of 600-800 ppm leads to death. The maximum concentration that workers can permanently be exposed to for 10 minutes without adverse health effects is 50 ppm. The entry of H_2_S gas into underground is one of the most important geological hazards. Tunneling in these areas requires special considerations. Hydrogen sulfide gas are released by many industries, including petrochemicals, agriculture, and sewage treatment in occupational and public environments. Addressing the risks and challenges related to the entry of the gas into tunnels is very difficult and expensive. One of the important tasks in this condition is to predict and estimate the risk of H_2_S gas in the underground areas and identify the appropriate method addressing engineering and environmental problems. There are several solutions for detecting hydrogen sulfide gas, including optical nanosensors, which are simple structured, portable, and cost-effective.

Metal nanoparticles are able to revolutionize sensor manufacturing due to their very small size, high specific surface area, electrical conductivity, and unique optical properties. Among the metal nanoparticles, silver has received much attention in nanosensor systems. Silver reacts with hydrogen sulfide favorably, and produces black precipitate of silver sulfide. This metal is mostly used in the form of nanoparticles in composites, since nano scaled particles have different properties compared to the mass state, and are interesting due to the increase in the ratio of surface area to volume leading to more surface reactivity. The detection mechanism is based on how silver nanoparticles affect the electrical properties of the nanocomposite, or change in the optical properties of the local surface plasmon resonance of the silver nanoparticles as a result of the reaction with gas.

The production of nanoparticles through biological method involves the use of different biological systems including simple prokaryotic organisms such as bacteria to complex Eukaryotes such as fungi and plants. Today, the commercial use of nanoparticles is rapidly advancing. The physical, chemical, and physiological properties of nanoparticles are influenced by their size and morphology. Bacteria, as biological nanofactories for the production of nanoparticles have received much attention. Researchers have found that microorganisms can be used as living factories for the synthesis of metal nanoparticles such as gold and silver nanoparticles. Furthermore, researchers prefer the biological synthesis method since control and distribution of the size of nanoparticles synthesized by this method is better than others. Streptomyces is a successful example used in the production of metal nanoparticles such as gold and silver nanoparticles.

Biosensors are tools with biological material base, and currently are widely used in various fields. These sensors are used as powerful tools to identify and sense biological molecules. Gas biosensors are widely used in detecting negligible amounts of toxic and flammable gases. Currently, one of the ways to detect H_2_S gas is to fabricate a hydrogen sulfide biosensor based on zinc oxide microstructures. ZnO rod structures can be synthesized on quartz substrates without the presence of a catalyst or buffer through the chemical vapor deposition (CVD) method in a horizontal cylindrical furnace.

In this research, advanced technologies such as biotechnology and nanotechnology has been used, which, in addition to high accuracy and sensitivity, enables the selective identification of a specific substance (hydrogen sulfide). Accordingly, a nano biosensor has been fabricated using silver nanoparticles for hydrogen sulfide gas detection. The method reviewed in this study is cost-effective and can replace the existing methods. It is worth noting that in most of the articles with silver as gas detector, the change of optical properties was considered. However, in this study, an attempt has been made to use a simple and visual method without the need for additional equipment for gas detection.

## II. Review of Literature

In recent years, studies have increased on the detection of hydrogen sulfide gas in low concentrations using biological structures. Characteristics such as simplicity, high sensitivity, low response time and affordability, have caused biosensors to be widely used in food analysis, environmental control, clinical diagnosis, pharmaceutical, and agricultural industries. Gas biosensors are widely used in detecting very low amounts of gases. The use of nanoparticles as a new innovation in detection technology leads to improvements in sensitivity, and multiple-target detection. Nanomaterial-based electrochemical biosensors possess most of the above-mentioned advantages such as high sensitivity, suitable selectivity, fast detection and cost-effectiveness; therefore, electrochemical sensors with high sensitivity and selectivity have been studied for many analytes. Researchers have found that microorganisms can be used as living factories for the synthesis of metal nanoparticles such as gold and silver.Moreover, researchers prefer the biological synthesis method since the control and distribution of the size of nanoparticles synthesized by this method is better than others.

Shadfar et al. conducted research entitled Measurement of Nitrate Concentration in Aqueous Media Using an Electrochemical Nanosensor Based on Silver Nanoparticles-Nanocellulose/Graphene Oxide, in 2017. In this research, researchers fabricated sensors based on nanocomposite consisting of nanocellulose and silver nanoparticles. The results showed that this method is highly precise and accurate, and the fabricated nanosensor with long-term stability is able to detect nitrate ions in aqueous solutions without the intervention of interfering factors [1].

In 2013, Soltaninejad et al. conducted research titled Extracellular Biological Production of Silver Nanoparticles by Streptomyces M67 Bacteria. Considering the compatibility of nanoparticles with the environment, they placed the mentioned bacterial colonies in a container containing silver nitrate. After 10 hours, the color of the solution started to change, and after 24 hours, the color of the solution changed from colorless to dark brown. They proved the production of silver nanoparticles by this species using spectrometry with the maximum absorption at 450 nm [2].

In 2016, Ghazizadeh et al. have conducted a research, Preparation of Gas Sensor Based on Polymer Nanocomposite for Qualitative Detection of Hydrogen Sulfide. In this research, they have used porous nanocomposite films of polyurethane, silver, and silver polyvinyl chloride containing 7 wt% of silver, as a qualitative hydrogen sulfide sensor. The results showed that when the samples were exposed to 50 ppm of hydrogen sulfide gas for 10 minutes, black spots appeared on the surface, and disappeared after exposure to air for 20 minutes. This type of sensors was suggested due to their portability and cost-effectiveness [3].

In 2014, Bermoz et al conducted a review article expressing the importance of soil Streptomyces and its prominent metabolites, concluding that Streptomyces can be used in nanobiotechnology studies and the production of nanoparticles of gold, silver, selenium, tellurium and other valuable elements [4].

Kashi et al conducted a research titled Microbial Synthesis of Silver Nanoparticles in 2018. They have investigated bacteria regarding their ability to regenerate heavy metal ions as biological factories for the production of nanoparticles. The researchers stated that with the bacterial production of nanoparticles as a green and one-step method, the problems and disadvantages of production through physical and chemical methods can be resolved. In this study, bacterial variants were extracted from the soil of Nakhlak Mine, and it was concluded that the bacteria extracted from this mine can be considered as a good biological source for the production of silver nanoparticles [5].

## III. Materials and Method

1. Hydrogen sulfide gas
2. Acetone
3. Gellan gum
4. Alginate
5. Ethylene dichloride
6. Silver nitrate
7. Broth culture medium
8. Nutrient agar culture medium
9. Streptomyces sp variant M67

### A. Preparing Pure Variants of Streptomyces sp Variant M67

Considering the possibility of extracellular production of silver nanoparticles in Streptomyces bacteria, and the resistance of this species to the antimicrobial properties of silver, the active culture of this bacterium was prepared in the Microbiology Laboratory of Shahid Beheshti University, Tehran, Iran.

### B. Biosynthesis of Silver Nanoparticles inside the Bacteria

To prepare nutrient agar culture medium, 2.8 gr of this powder was added to 100 ml of distilled water, dissolved by heat, and sterilized inside the autoclave, then distributed in the plates.

The bacterial variant was cultured linearly in nutrient agar culture medium under sterile condition. After 24 hours of preserving the plate warm in the incubator, bacterial colonies were formed.

Then, the colonies grown in the plate were transferred to an Erlenmeyer flask containing 200 ml of Nutrient Broth culture medium, and incubated in a shaker incubator for 24 hours at a speed of 200 rpm at a temperature of 30° C.

After 24 hours of cell growth and proliferation, the bacteria were centrifuged at 4000 rpm for 25 minutes, separated from the culture medium.

Further, the cells were washed 3 times with physiological serum. 0.5 gr of the wet weight of cell masses was removed and placed in 50 ml of 3.5 mM silver nitrate solution with pH 7 in a 250 ml Erlenmeyer flask in the dark for 48-72 hours, under sterile conditions, incubated in a shaker incubator at a temperature of 30° C at a speed of 200 rpm.

### c. Visible and Ultraviolet Spectrophotometry (UV-Vis)

Silver nanoparticles absorb electromagnetic waves in 400-470 nm. Due to the extracellular production of silver nanoparticles, 50 µl of the isolated solution was poured into the cuvette and 25 µl of distilled water was added to it, and spectroscopy was carried out in the range of 390-470 nm.

### D. Microencapsulation of Bacteria in Biocomposite

The bacteria were microencapsulation using gellan gum and alginate. In this step, 1 wt% solutions of alginate and gellan were prepared, and combined with a ratio of gellan to alginate of 3:7, mixed for one hour at room temperature on an Erlen shaker at 150 rpm.

The solid of bacterial cells isolated through centrifugation was washed twice with sterile normal saline, and added to the prepared gellan and alginate solution along with 10 µl of the centrifuged solution. The resulting solution was placed in a Bain-Marie at a temperature of 60° C.

After that, the solution was mixed with 20 ml of 10% aqueous acetone solution containing 50 µg of EDC (ethylene dichloride) on a heater stirrer at a temperature of 60° C, then placed in the oven at 60° C for 12 hours.

In the coating stage, glass pieces with a size of 24 mm*24 mm were used. First, they were washed well with soap and water and placed in the oven. Then, the prepared solution was applied on these pieces through deep-coating method.

### E. Determining the Viscosity and Porosity of Biocomposite

In order to determine the viscosity of the biocomposite solution, 50 ml of the prepared solution was poured into the vessel of the viscometer, and after the temperature of the sample reached 25°, the dynamic viscosity was measured. Moreover, the porosity of the membrane was examined using a microscopic image.

### F. Validating the Performance of the Detector

In this test, the biocomposite was placed inside a 10 cm*10 cm*10 cm plexiglass container, connected to a cylinder of hydrogen sulfide gas diluted with nitrogen on one side, and to the open air on the other side. After establishing the gas flow in different time periods, the color of the sensor changed by time. It is worth noting that the samples were tested in two forms, with biocomposite substrates encapsulated with bacteria (test sample) and without bacteria (control sample). The synthetized biocomposite was suspended in the middle of the cube and exposed to hydrogen sulfide gas. The air quality inside the container was examined using a digital thermometer and hygrometer.

## III. Results and Discussion

### A. Confirming the Synthesis of Silver Nanoparticles in Bacteria

The formation of bacterial colonies, their resistance to the antibacterial property of silver nitrate, and the color change of solution obtained from the linear culture of bacteria from pale yellow to brown (see Figure 1) proved that the bacteria is capable of producing silver nanoparticles.

**Figure 1.**
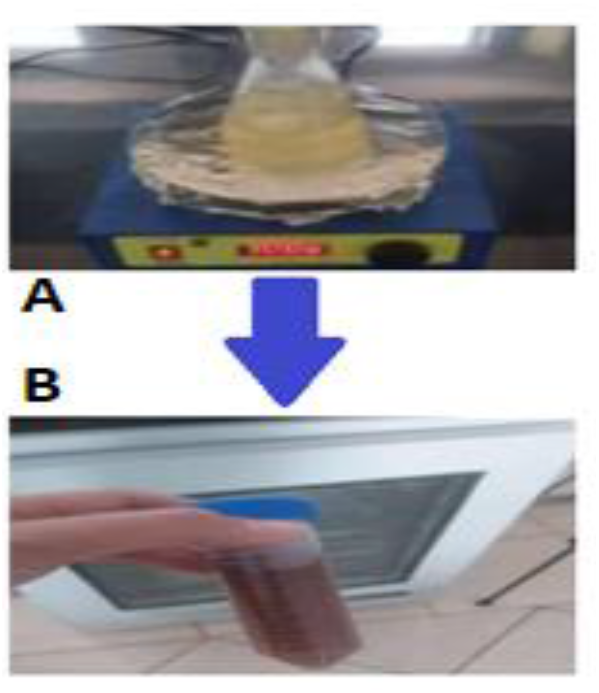
A) The synthesis of nanoparticles in the microbial environment. The color of the solution is yellow at the beginning of the synthesis process. B) The color change of the solution to brown indicates the synthesized nanoparticles in the solution.

Since the silver nanoparticles show absorption spectrum in the range of visible and ultraviolet wavelengths, the UV-Vis spectroscopy method was used for confirmation (see Figure 2). The maximum absorption is observed at the wavelength of 420 nm.

**Figure 2.**
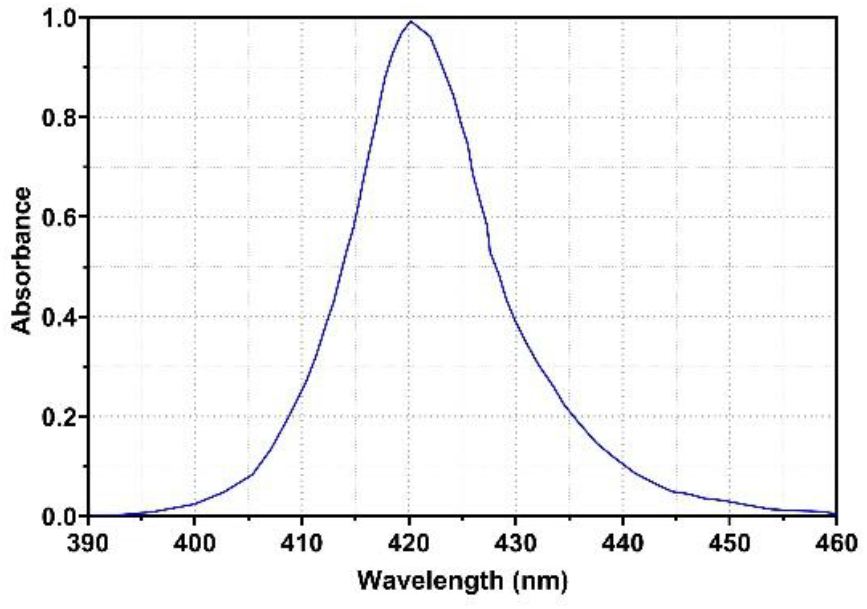
UV-Vis spectroscopy analysis of silver nanoparticles

Based on the wavelength of the observed peak in Figure 2, and the data presented in the graph [6] in Figure 3, since the peak is slightly broadened, the size of the nanoparticles can be reported as 50 nm, with an acceptable approximation.

**Figure 3.**
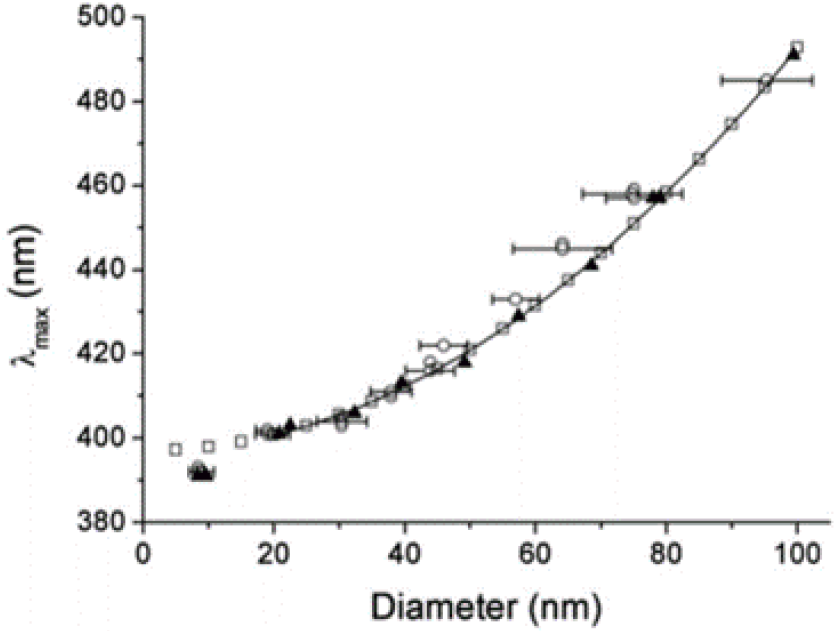
The experimental (hollow circles) and simulated (hollow squares) absorption peak of silver nanoparticles

### A. Investigating the Porosity and Viscosity of Biocomposite

The porosity and viscosity of the biocomposite were investigated with the aim of measuring the gas permeability in biocomposite. These two parameters directly affect the response time of the detector.

The porosimetry analysis based on optical microscope (Figure 4), and viscosity measurement were conducted, the obtained figures reported in Table 1 are considered acceptable compared to other studies conducted in this field. It is found that the efficiency of the detector is not limiting in this section.

**Table 1.** The results of porosimetry analysis and viscosity measurements

| Material | Dynamic Viscosity | Porosity Percentage |
| --- | --- | --- |
| Gellan alginate biocomposite containing silver nanoparticles | 1.09 | 63.58 |

**Figure 4.**
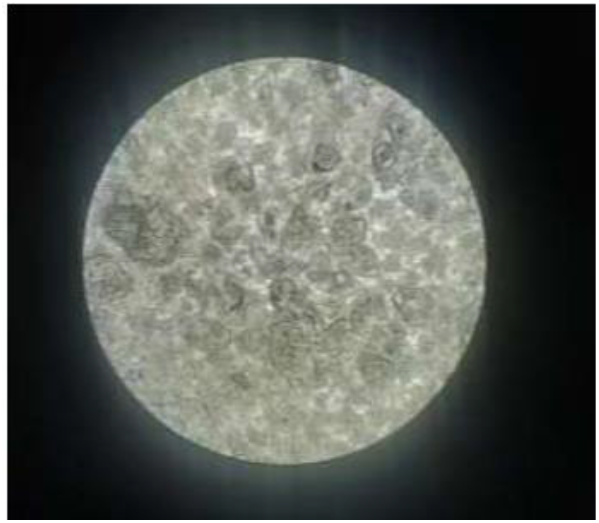
The porous structure of the biocomposite

### B. Investigating the Performance of Hydrogen Sulfide Gas Detector

Based on the observations, hydrogen sulfide, a gas with weak acid properties, is absorbed by the biocomposite substrate, reacts with silver nanoparticles produced by bacteria, and produces black precipitate of silver sulfide, according to equation 1.

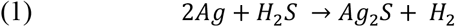

In order to investigate the performance of the detector, the input of H_2_S gas was prepared in 5 concentrations of 10, 20, 30, 40 and 50 mg/m3, and at different concentrations, the first time when the color of the biocomposite changed was recorded. The experiment was repeated 3 times on different days. The temperature and humidity were preserved unchanged, measured using a digital thermometer and hygrometer (see table 2).

**Table 2.** The figures related to the performance of H_2_S detector

| The input gas concentration | The average of temperature | The average of humidity | The time when the color started to change |
| --- | --- | --- | --- |
| 10 | 28.6° C | 79.32% | Unchanged |
| 20 | 28.6° C | 79.32% | Slightly changed |
| 30 | 28.6° C | 79.32% | 18 min |
| 40 | 28.6° C | 79.32% | 15 min |
| 50 | 28.6° C | 79.32% | 12 min |

Figure 5 shows the color change of the silver nanoparticles of the test sample compared to the control sample, confirming the performance of the detector.

**Figure 5.**
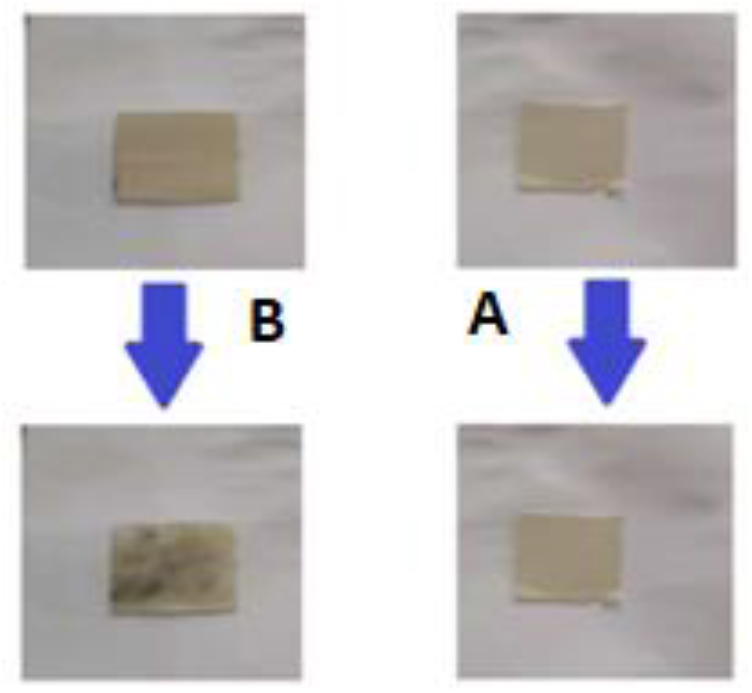
A) The color of the control sample, which does not contain silver nanoparticles, did not change when exposed to 50 mg/l hydrogen sulfide gas. B) The test sample with silver nanoparticles, goes through a color change when exposed to 50 mg/l hydrogen sulfide gas.

## V. Conclusion

According to the findings of the research on using Streptomyces bacteria as a biofactory to produce silver nanoparticles, the research has fulfilled acceptable results. Compared to the work of Dehghani et al.[7], Colorimetric biosensors based on silver nanoparticles, which is basically similar to the biosensor fabricated in the mentioned research, was accomplished through a green and sustainable method. Moreover, this biodetector is more simple-structured and cost-effective, which is how it is more advantageous than that of the Dehghani et al.

The working principle of the nanobiodetector is colorimetric method, a simple method interpreting the results with no need for a special device. Compared to the biosensor fabricated by Ghazizadeh et al.[3], in which a polymer nanocomposite-based gas sensor was used for the qualitative detection of hydrogen sulfide, the type of materials used in the biocomposite of the present study is more cost-effective, and sensitive. The nanocomposite of the so-called research was fabricated using PVS and PU, where the nanocomposite of this study is fabricated using Gellan gum and alginate, which are more accessible, cost-effective, and acceptably sensitive with fine response time.

